# Alginate-Coated Collagen Hydrogel Tubes for Scalable Cell and Biotherapeutic Particle Manufacturing

**DOI:** 10.64898/2026.05.29.728820

**Authors:** Xinran Wu, Ying Pan, Yakun Yang, Li Han, Yong Wang, Aziza P. Manceur, Yuguo Lei

## Abstract

Efficient, scalable, and cost-effective production of mammalian cells and biotherapeutic particles remains a major challenge for both research and clinical applications. Conventional 2D and 3D culture systems suffer from low volumetric yields, poor scalability, and high costs. Previously, we developed collagen hydrogel tube microbioreactors (ColTubes) that support high-density, high-viability cell culture by preventing excessive cell aggregation and minimizing hydrodynamic stress. However, ColTubes exhibit adhesion to culture vessels and to each other, and leaked cells frequently attach to outer tube surfaces — behaviors that would limit scalability. Here we introduce AlgColTubes: collagen hydrogel tubes coated with a thin, ionically crosslinked alginate layer to overcome these limitations. Scanning electron microscopy confirms that alginate penetrates the collagen wall and forms a stable interpenetrating hydrogel network, whose depth can be tuned by coating concentration and duration. The alginate coating remains structurally intact under static and dynamic culture conditions without impairing nutrient transport or cell growth. AlgColTubes eliminate tube-tube and tube-vessel adhesion, and prevent exogenous cell attachment to the outer surface, while maintaining cell viability and proliferation comparable to uncoated ColTubes. Their unique architecture — an adhesive collagen interior and non-adhesive alginate exterior — further enables a truncated-tube format for continuous release of biotherapeutic particles through open tube ends. We demonstrate that lentivirus is released from truncated AlgColTubes in a segment length-dependent manner, reaching ~100% release efficiency at 1-mm segment lengths. AlgColTubes provide a scalable, cost-effective platform for high-yield cell and particle manufacturing, with broad potential across basic research, translational studies, and industrial bioprocessing.

## A. Introduction

Large-scale production of mammalian cells and biotherapeutic particles is constrained by persistent limitations in current manufacturing platforms. Two-dimensional (2D) flask-based culture systems remain the most widely used laboratory tools, yet they provide microenvironments that differ fundamentally from the three-dimensional (3D) niches cells occupy in vivo — lacking structural extracellular matrix (ECM) support, cell-cell contact geometry, and the physiological gradients that govern growth and differentiation^1–3^. They have low productivity, high reagent and labor costs, and poor reproducibility and scalability^1–3^. Three-dimensional suspension cultures such as stirred-tank bioreactors were developed to address these limitations, but introduce new challenges: uncontrolled cell aggregation generates clusters that exceed the diffusion limit, creating hypoxic and nutrient-depleted cores that drive apoptosis and undesired differentiation, while the agitation simultaneously imposes damaging hydrodynamic shear^4,5^. These competing constraints result in low volumetric yields and batch-to-batch variability that worsens with scale-up, underscoring the need for fundamentally different culture architectures^1–3^.

Hydrogel tube microbioreactors represent a promising solution for creating cell-friendly, scalable culture environments^4–13^. In these systems, cells are encapsulated within microscale hydrogel tubes whose wall thickness and diameter can be controlled to keep all cells within the diffusion limit, ensuring efficient nutrient and waste transport throughout the culture^4^. Nanoscale pores in the hydrogel wall permit free passage of nutrients and growth factors while physically shielding cells from hydrodynamic stress. The tubular geometry provides continuous space for cell expansion and cell-cell interaction — critical for maintaining viability, proliferative capacity, and phenotypic integrity during large-scale culture^4,5^. In our prior work, we demonstrated that alginate-based hydrogel tubes (**AlgTubes**) support volumetric yields up to 5×10^8^ cells/mL — more than 200-fold greater than conventional stirred-tank bioreactors — across multiple cell types including hPSCs, endothelial cells, neural stem cells, vascular smooth muscle cells, and T cells ^4–13^.

To better replicate the native cellular microenvironment, we subsequently developed collagen hydrogel tube microbioreactors (**ColTubes**) using collagen — the primary structural protein of human connective tissues^14^. ColTubes provide integrin-binding sites and native fibrillar nanoarchitecture that alginate cannot offer, and they support high-density, high-viability cultures comparable to AlgTubes^14^. However, ColTubes adhere to culture vessels and to each other, and cells that leak from tube ends attach to outer surfaces and promote tube aggregation ^14^. These behaviors would impede suspension handling and throughput in any large-scale manufacturing context.

In this work, we report a simple and effective strategy to resolve this limitation: coating the outer surface of ColTubes with a thin, ionically crosslinked alginate hydrogel to create AlgColTubes. Alginate’s dense hydration shell and negative surface charge confer well-characterized non-fouling and anti-adhesive properties that prevent protein adsorption and cell attachment. Importantly, alginate diffuses into the collagen wall before crosslinking, forming a physically entangled interpenetrating network (IPN) rather than a loosely adsorbed surface layer — a mechanism that provides durable coating stability. We characterize the nanostructure, coating stability, cytocompatibility, and anti-adhesion performance of AlgColTubes under both static and dynamic conditions, and demonstrate a truncated-tube format for continuous lentiviral vector production. AlgColTubes thus combine the biological fidelity of collagen with the scalability-enabling surface chemistry of alginate, offering a versatile and readily translatable platform for next-generation cell and particle biomanufacturing.

## B. Materials and Methods

### Processing ColTubes

Commercial rat tail collagen I (Advanced BioMatrix, #5153) was used to prepare ColTubes. Collagen from other species, such as bovine, has also been tested for ColTube preparation with comparable success. A custom micro-extruder was designed using Fusion 360 (Autodesk) and fabricated using a stereolithography 3D printer (Form 3B+, Formlabs) for collagen tube production. Three inlets were connected to syringes mounted on syringe pumps to precisely control flow rates. A syringe containing a 1.5% methylcellulose or hyaluronic acid solution with single cells was connected to the core flow channel. A pre-chilled syringe loaded with collagen solution was placed in a custom ice box, connected to the shell flow channel. A syringe containing 50 mM HEPES buffer was attached to the sheath flow channel. The outlet of the micro-extruder was submerged in 37°C HEPES buffer maintained using a heating pad. Once activated, collagen tubes were continuously generated and collected into HEPES buffer. The HEPES buffer was subsequently replaced with pre-heated cell culture medium.

### Coating ColTubes with Alginate

ColTubes were immersed in alginate solutions at varying concentrations and for different durations according to the experimental design. Immediately afterward, the alginate-covered ColTubes were gently transferred into 100 mM CaCl_2_ solution to induce ionic crosslinking and form a stable external alginate shell. After crosslinking, AlgColTubes were rinsed in culture medium to remove excess CaCl_2_.

### Confocal microscopy imaging

To visualize the nanostructures of collagen and alginate in AlgColTubes, Alginate-Fluorescein (HAWORKS, AL-FITC-LV) was used to visualize the alginate layer, and ATTO 594 NHS ester (ATT Bioquest, 2860) was used to label collagen fibers per the manufacturer’s instructions. A 10 mM stock of ATTO 594 NHS ester was diluted in PBS to 1 µM working concentration. ColTubes were first coated with Alginate-FITC as described above. The resulting AlgColTubes were incubated with 1 µM ATTO 594 NHS ester for 15 min at room temperature, followed by two PBS washes to remove unbound dye. Confocal fluorescence images were acquired using an Olympus FV3000 microscope with a 10× objective.

### Scanning Electron Microscopy (SEM)

Nanostructures of hydrogel samples were analyzed using a Zeiss SIGMA VP-FESEM. Samples were dehydrated through a graded ethanol series (25%, 50%, 70%, 85%, 95%, 100%), followed by critical point drying (Leica EM CPD300). Samples were sputter-coated with a 7 nm iridium layer (Leica EM ACE600) to enhance conductivity. SEM images were acquired under high vacuum at 5 kV accelerating voltage, 5 mm working distance, at magnifications from ×50 to ×5,000.

### Culturing hPSCs in AlgColTubes

H9 human embryonic stem cells (WiCell, WA09) were used for this study. All experiments involving human embryonic stem cells were performed in strict accordance with institutional guidelines and approved by the appropriate ethics committee at Pennsylvania State University. The H9 hPSCs was obtained from the WiCell Research Institute and authorized for research use. All procedures complied with applicable national regulations and policies governing the ethical use of human stem cells in research.

For a typical culture, 10 µL of cell-laden AlgColTubes was suspended in 2 mL E8 medium supplemented with 10 µM Y-27632 in a 6-well plate, and maintained at 37°C, 5% CO_2_, 21% O_2_. Medium was replaced daily. To passage cells, medium was removed, and AlgColTubes were dissolved by incubating with 0.5 mM EDTA and 0.2 mg/mL Collagenase P for 15 minutes. The cell mass was collected by centrifugation at 100 × g for 5 minutes and treated with Accutase at 37°C for 10 minutes to yield single-cell suspensions for re-culture or cryopreservation.

### Culturing HEK 293SF-LVP #3E9 cells and lentivirus production in truncated AlgColTubes

The HEK 293SF-LVP #3E9 cell line was obtained from the National Research Council Canada^15^. 5 µL of cell-laden AlgColTubes was suspended in 2 mL Hycell-Transfx-H medium supplemented with 4 mM L-Glutamine and 0.1% Poloxamer 188; medium was changed daily. After 5 days of culture, lentiviral expression was induced by adding 80 µg/mL cumate and 10 mM coumermycin; 7 mM sodium butyrate was added at 18 hours post-induction to enhance lentiviral production. Following induction, AlgColTubes were cut into segments of 3 cm, 1 cm, or 1 mm. At 72 hours post-induction, conditioned medium containing secreted lentivirus was collected and titrated via infecting HEK293 cells. For positive controls, AlgColTubes were fully digested to release all retained lentivirus prior to titration.

### Statistical analysis

All data were analyzed using GraphPad Prism 8 and are presented as mean ± SEM. Statistical comparisons used one-way ANOVA for three or more groups, log-rank test for survival analyses, or unpaired two-tailed t-tests for two-group comparisons. Significance is denoted as *: p<0.05, **: p<0.01, ***: p<0.001.

## C. Results

### Fabrication and Structural Characterization of AlgColTubes

AlgColTubes were fabricated by a sequential two-step process (Fig. 1A). ColTubes were first produced by coaxial microextrusion of a cell-laden methylcellulose core, collagen shell, and HEPES buffer sheath into heated HEPES buffer, inducing rapid gelation of the collagen shell through simultaneous pH neutralization and temperature increase^14^. The resulting ColTubes were then immersed in alginate solution and transferred to 100 mM CaCl_2_ for ionic crosslinking of the alginate coating. The detailed ColTube fabrication method is described in our prior publication^14^ and in Fig. S1. Confocal imaging of dual-labeled tubes (ATTO 594–collagen; FITC–alginate) confirmed that alginate coating thickness increases monotonically with both dipping time (5–60 min at 1.5% alginate; Fig. 1B) and alginate concentration (at fixed 15 min dipping time; Fig. 1C), providing independent and predictable control over the coating layer.

**Fig. 1.**
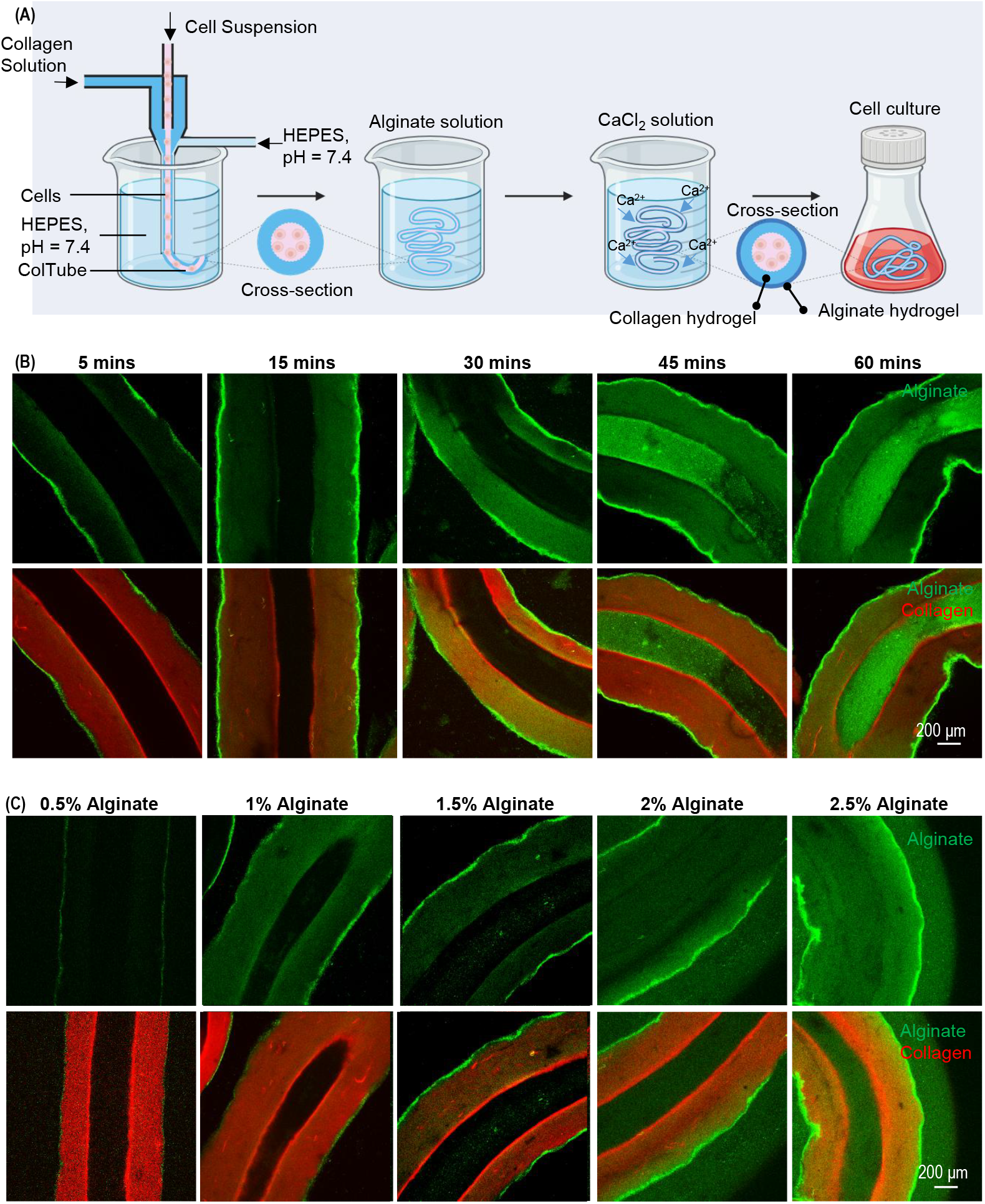
Coating ColTubes with Alginate Hydrogel. (A) Schematic illustration of fabricating alginate-coated collagen hydrogel tubes (AlgColTubes). ColTubes are produced using a three-flow micro-extruder, then dipped in an alginate solution followed by immersion in a Ca^2+^ buffer to form a thin alginate hydrogel coating. (B) Coating thickness increases with longer coating times, while the alginate solution concentration is fixed at 1.5%. (C) Coating thickness increases with higher alginate concentrations, while the coating time is fixed at 15 min. In (B) and (C), collagen is labeled with a red fluorescent dye and alginate with a green fluorescent dye. Confocal imaging was used to visualize the coating.

SEM imaging revealed distinct nanoarchitectures for each tube type. AlgTubes exhibited a dense, isotropic mesh-like network throughout the wall, whereas ColTubes showed open collagen nanofiber bundles with large inter-fiber spaces (Figs. 2A, 2B). AlgColTubes displayed spatially graded hybrid structures that varied with coating parameters: the outer surface was dominated by a continuous, dense alginate mesh resembling AlgTubes, transitioning progressively to a collagen fibrillar interior toward the tube lumen (Fig. 2C–E). At 0.5% alginate for 15 min, alginate penetration was confined to the outer wall (ROI 1–2), while the middle and inner regions retained the native collagen nanofiber structure (ROI 3–4; Fig. 2C). Increasing alginate concentration to 2.5% produced full-thickness alginate penetration across all four wall regions (Fig. 2E). Similarly, extending coating time from 5 to 30 min at 1.5% alginate progressed from partial to complete wall penetration (Fig. 3C–E). These results demonstrate that alginate diffuses into and crosslinks within the collagen matrix, forming a stable interpenetrating hydrogel network whose depth is tunable by adjusting coating concentration and duration.

**Fig. 2.**
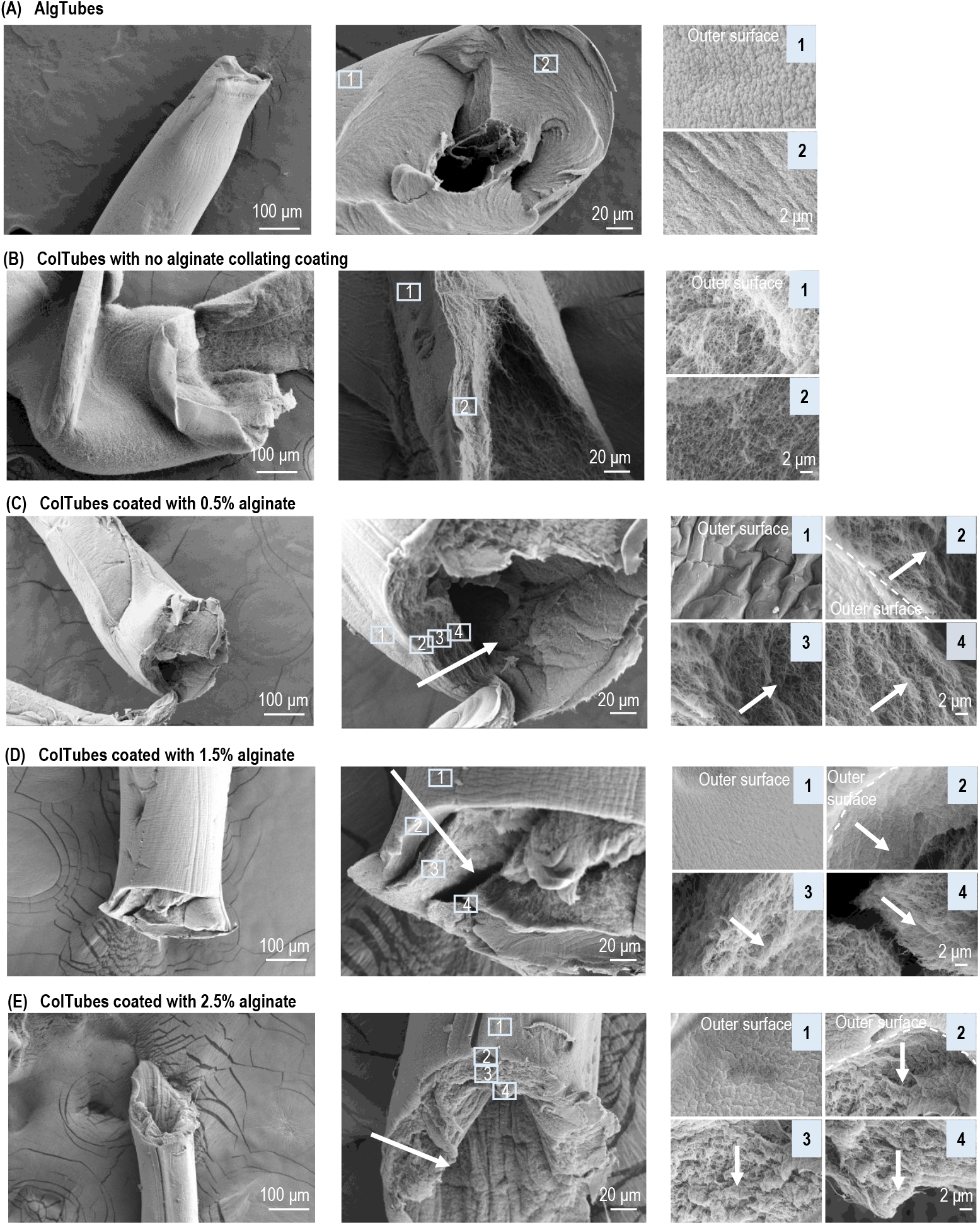
Nanostructure of ColTubes coated with different concentrations of alginate hydrogel. SEM images of AlgTubes (A), uncoated ColTubes (B), and ColTubes coated with 0.5% (C), 1.5% (D), or 2.5% (E) alginate hydrogels for 15 min. Low magnification SEM images show the overall tube morphology for each condition. High magnification SEM images corresponding to ROI 1–4 reveal progressive changes in tube wall texture toward the lumen. A continuous alginate layer formed on the outer surface (ROI 1), and alginate gradually diffused from the outer surface toward the inner surface of the ColTube (ROI 1–4). White arrows indicate the direction of alginate diffusion.

**Fig. 3.**
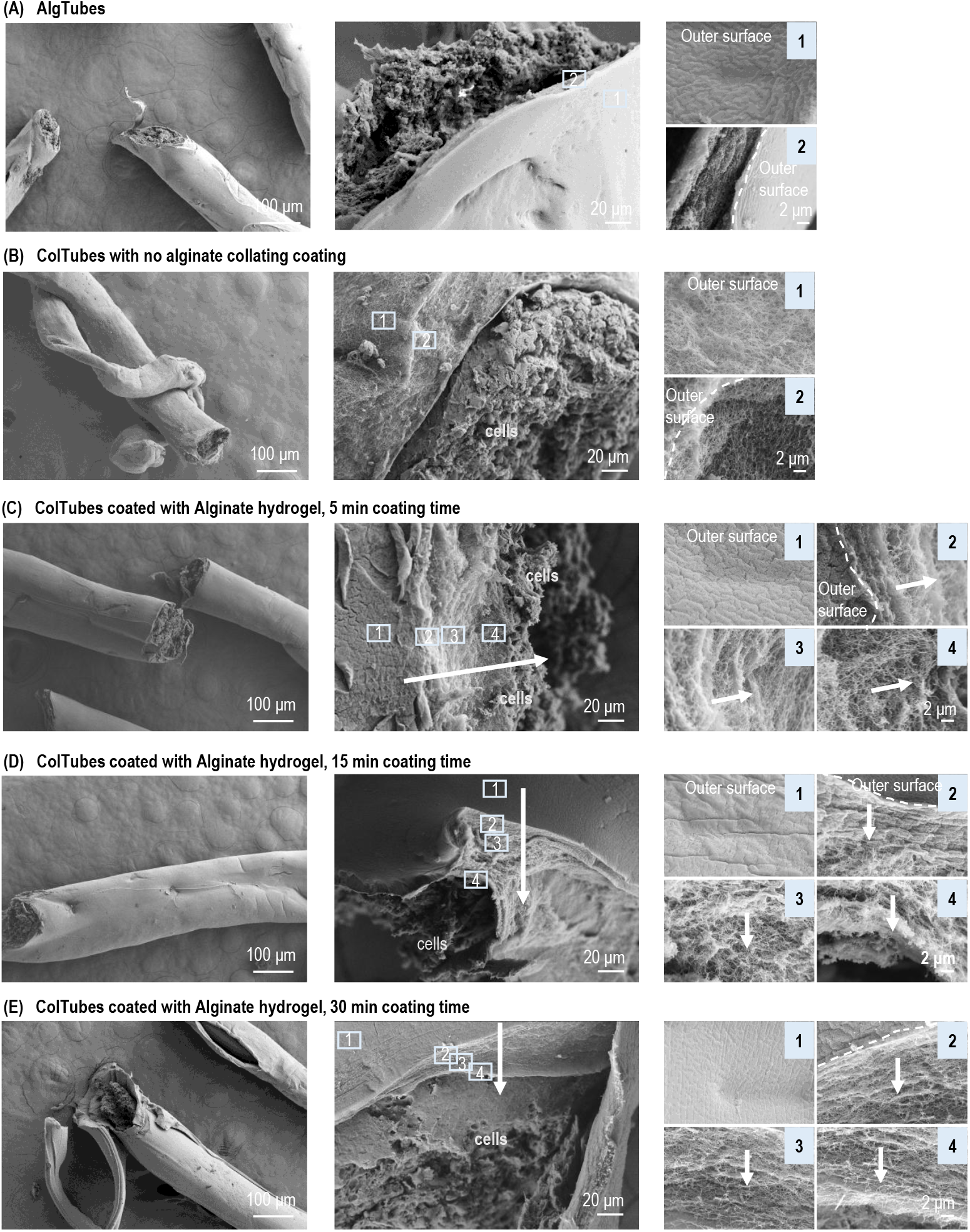
Nanostructure of AlgColTubes with different alginate coating durations. hPSCs were cultured in AlgTubes (A), uncoated ColTubes (B) or AlgColTubes generated using a constant 1.5% alginate solution and coated for 5 min (C), 15 min (D), and 30 min (E). Low magnification SEM images show the overall tube morphology under each condition. High magnification SEM images corresponding to ROI 1–4 reveal progressive changes in tube wall texture toward the lumen. A continuous alginate layer formed on the outer surface (ROI 1), and alginate gradually diffused from the outer surface toward the inner surface of the ColTube (ROI 1–4). White arrows indicate the direction of alginate diffusion.

### Coating Stability and Cytocompatibility

To assess whether the alginate coating is stable during culture and whether it affects cell behavior, hPSCs were loaded into ColTubes and AlgColTubes (5, 15, and 30 min coating times) and monitored for 5 days under static conditions using FITC-labeled alginate and dead-cell staining (Fig. 4). Fluorescence imaging revealed no detectable change in coating thickness or distribution across the 5-day observation period, confirming structural stability against medium exchange and metabolic conditions. Cell proliferation and viability were indistinguishable across all tube formats, with rare dead cells observed in all conditions — indicating that the alginate coating does not restrict nutrient or growth factor transport, and is non-cytotoxic at all coating times tested. In industrial cell manufacturing, mixing by stirring or orbital shaking is essential for medium homogenization. We therefore repeated these experiments under orbital shaking at 60 rpm (Figs. 5, 6). Results were virtually identical to static culture: the alginate coating remained intact, cell growth was robust, and dead cells were rare across all AlgColTube conditions (Figs. 6, 7). These results collectively confirm that the alginate IPN coating is mechanically stable under both static and dynamic culture conditions without compromising mass transport or cell health.

**Fig. 4.**
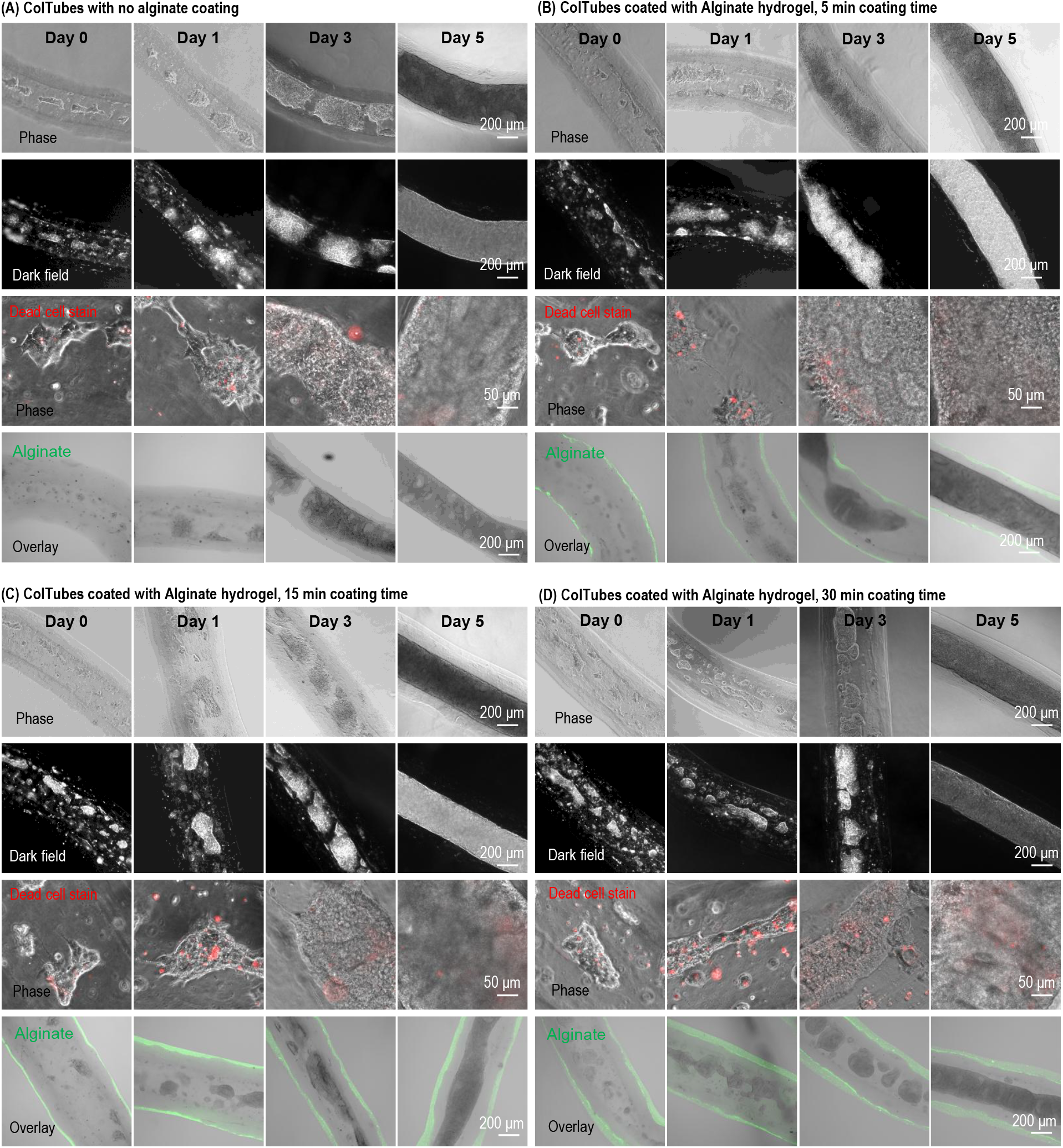
Stability and cytocompatibility of alginate hydrogel coating in static culture. hPSCs were cultured in uncoated ColTubes (A) or AlgColTubes prepared using a 1.5% alginate solution and coated for 5 min (B), 15 min (C), or 30 min (D). hPSCs exhibited healthy growth in all conditions with minimal cell death, as indicated by dead cell staining. The alginate hydrogel coating remained stable throughout the culture period. Images include phase contrast, dark-field, and phase overlays with alginate fluorescence. Cells are cultured statically without shaking.

**Fig. 5.**
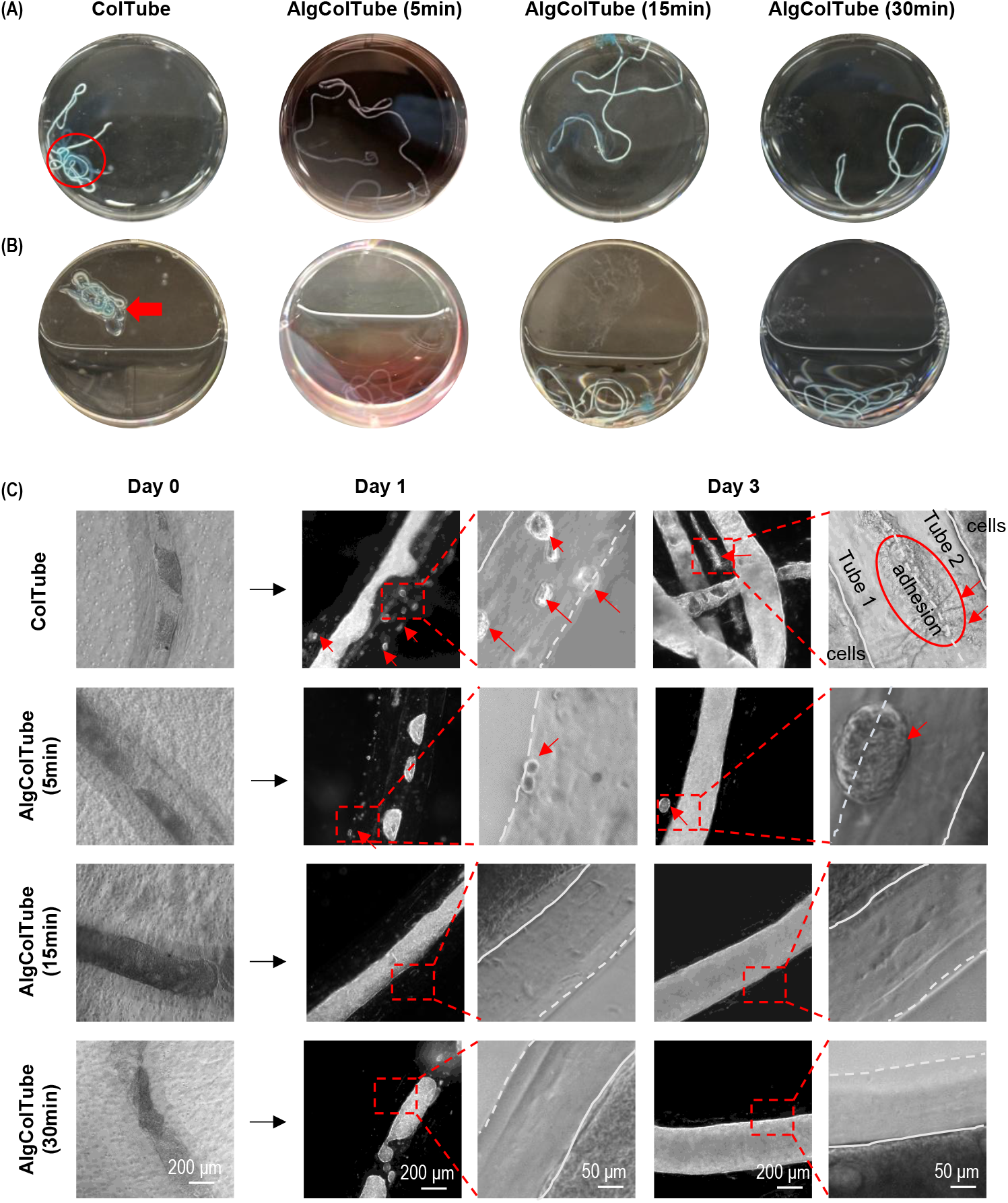
Alginate coating reduces surface adhesion of ColTubes under static conditions. (A) ColTubes and AlgColTubes were suspended in PBS. ColTubes adhered to neighboring tubes (red circle), whereas AlgColTubes did not. (B) When the plate was tilted, AlgColTubes settled freely, while ColTubes frequently adhered to the plate bottom (red arrow), indicating strong interaction with the culture vessel. (C) ColTubes and AlgColTubes containing hPSCs were cultured in a 6-well plate. Additionally, single hPSCs were added to the medium to assess adhesion to tube surfaces. Cells adhered to the outer surface of ColTubes (red arrows). A 5-min alginate coating reduced adhesion, while 15-min and 30-min coatings completely eliminated cell attachment. Cells are cultured statically without shaking. Solid and dashed white lines denote the inner and outer walls of ColTube or AlgColTube, respectively.

**Fig. 6.**
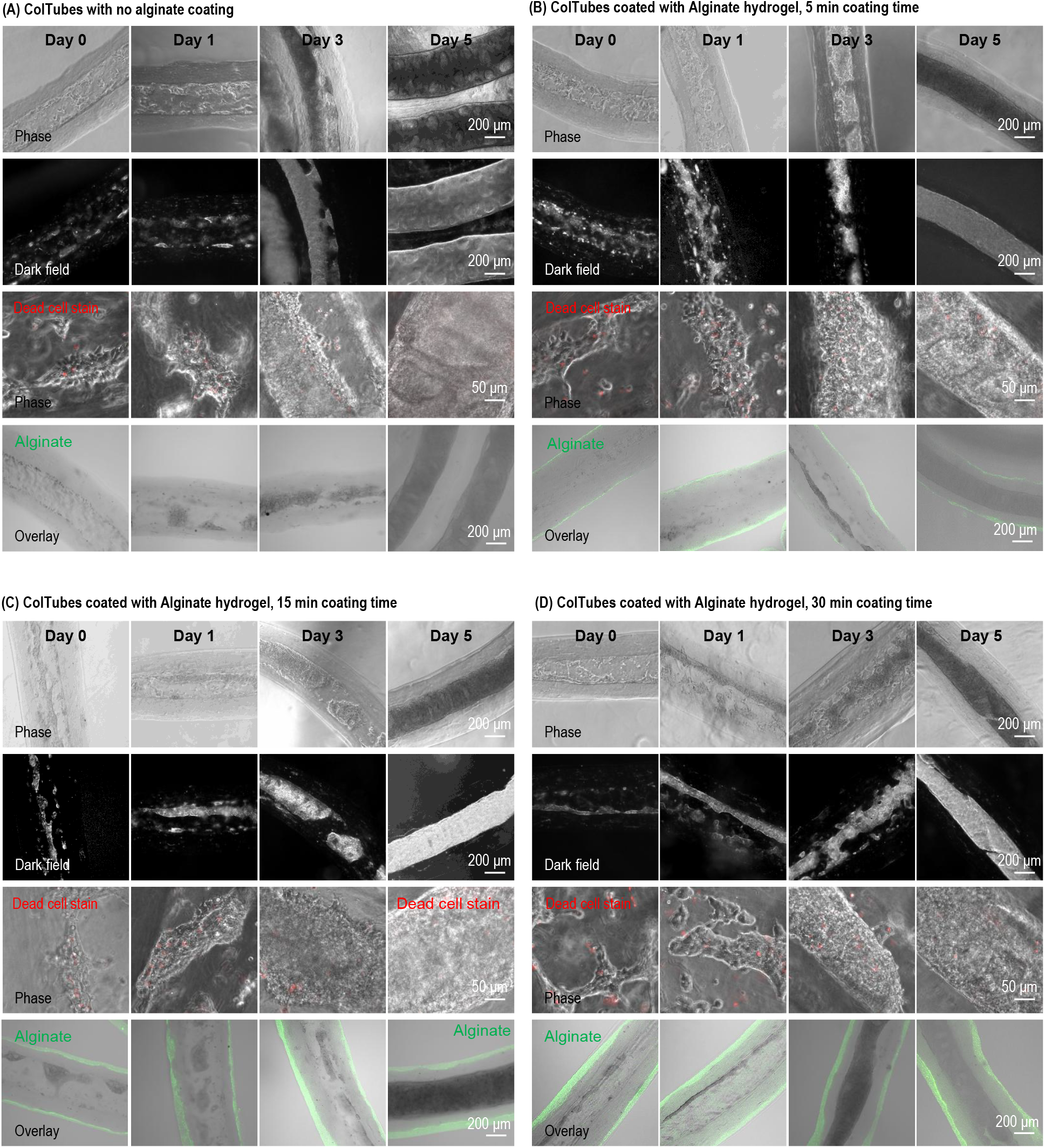
Stability and cytocompatibility of alginate hydrogel coating in dynamic culture. Human pluripotent stem cells (hPSCs) were cultured in uncoated ColTubes (A) or AlgColTubes fabricated with 1.5% alginate and coated for 5 min (B), 15 min (C), or 30 min (D). hPSCs exhibited healthy growth in all conditions with minimal cell death, as indicated by dead cell staining. The alginate hydrogel coating remained stable throughout the culture period. Images include phase contrast, dark-field, and phase overlays with alginate fluorescence. Cells were cultured dynamic in a shaking incubator with 60 rpm.

**Fig. 7.**
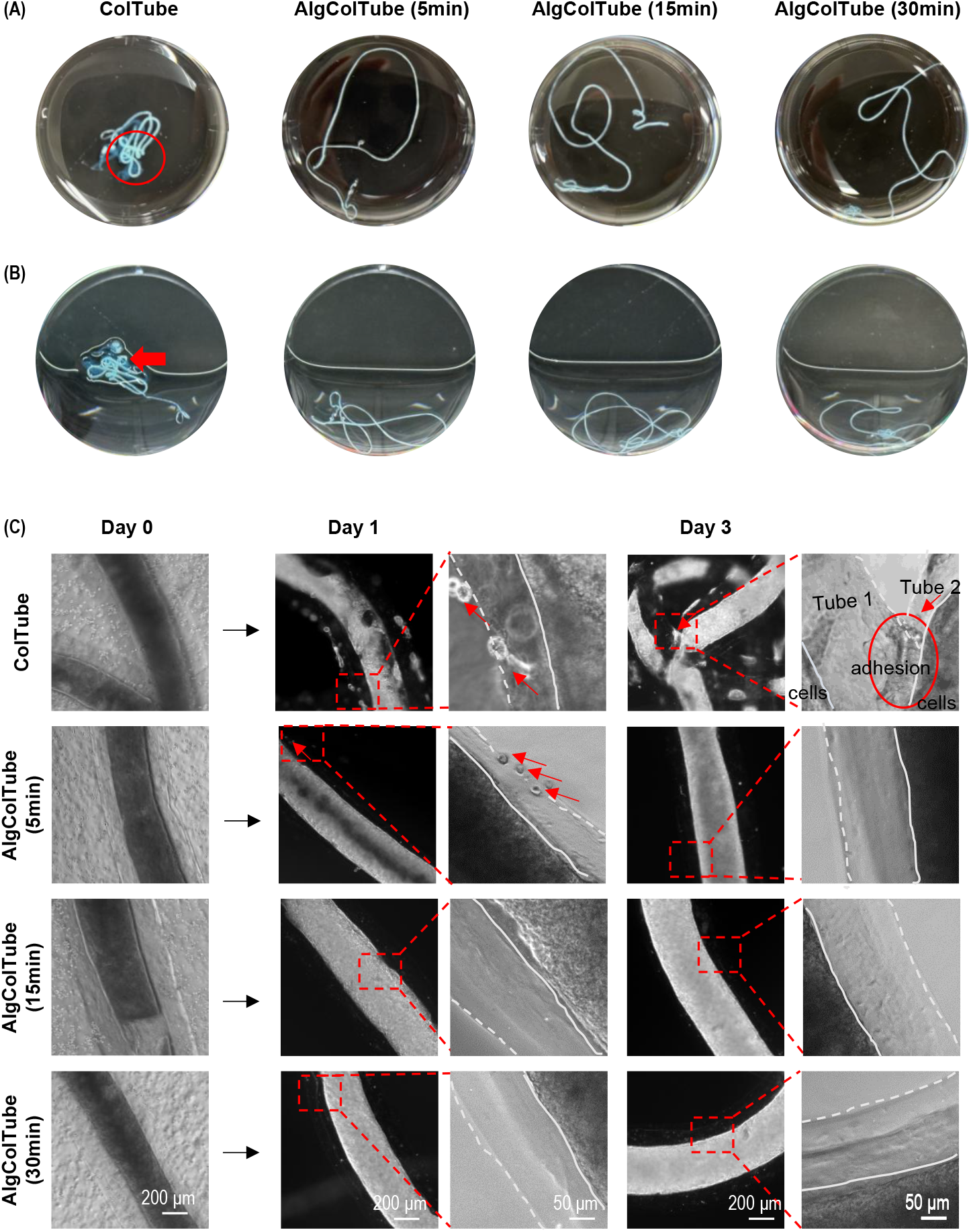
Alginate coating reduces surface adhesion of ColTubes under dynamic conditions. (A) ColTubes and AlgColTubes were suspended in PBS and shaken at 60 rpm. ColTubes adhered to neighboring tubes (red circles), whereas AlgColTubes remained non-adherent. (B) When the plate was tilted, AlgColTubes settled freely, while ColTubes frequently adhered to the plate bottom, indicating strong interaction with the culture vessel. (C) ColTubes and AlgColTubes containing hPSCs were cultured in a 6-well plate with orbital shaking at 60 rpm. Single hPSCs were added to the medium to evaluate adhesion to the tube surface. Cell attachment was observed on the outer surface of ColTubes (red arrows). A 5-min alginate coating partially reduced cell adhesion, while 15-min and 30-min coatings effectively eliminated cell attachment. Solid and dashed white lines denote the inner and outer walls of Coltube or AlgColTube, respectively.

### AlgColTubes Eliminate Surface Adhesion Under Static and Dynamic Conditions

A defining limitation of uncoated ColTubes is collagen’s inherent stickiness, which promotes tube-tube aggregation, tube-vessel adhesion, and the colonization of outer surfaces by leaked cells. To systematically evaluate the impact of alginate coating on these behaviors, we conducted a series of adhesion assays under both static and dynamic conditions.

In PBS suspension, uncoated ColTubes formed inter-tube aggregates and adhered to the plate bottom upon tilting, while AlgColTubes settled freely with no observable adhesion at either static (Fig. 5A, B) or 60-rpm dynamic conditions (Fig. 7A, B). To assess outer-surface cell fouling, single hPSCs were introduced into cultures containing ColTubes or AlgColTubes loaded with cells, and adhesion to outer surfaces was monitored over 3 days. hPSCs adhered robustly to ColTube outer surfaces; 5-min AlgColTubes showed partial reduction, while 15-min and 30-min coatings completely abolished outer-surface cell attachment under both static (Fig. 5C) and dynamic conditions (Fig. 7C). The anti-fouling effect of the alginate coating is attributable to its hydrophilic, negatively charged surface, which resists protein adsorption and cell attachment through a combination of steric and electrostatic repulsion — mechanisms well established for polyanionic hydrogel coatings. These results indicate that a coating time of ≥15 min at 1.5% alginate is sufficient to completely prevent all measured surface adhesion phenomena under physiologically relevant culture conditions.

### Truncated AlgColTubes Enable Continuous Biotherapeutic Particle Release

In intact long tubes, biotherapeutic particles produced by encapsulated cells, such as virus, cannot readily traverse the nanoporous hydrogel wall and are therefore trapped within the tube lumen, precluding continuous harvest without tube dissolution (Fig. 8). We hypothesized that cutting long AlgColTubes into short segments with open ends would allow particles to exit through the exposed ends into the surrounding medium, enabling continuous production and collection (Fig. 8A).

**Fig. 8.**
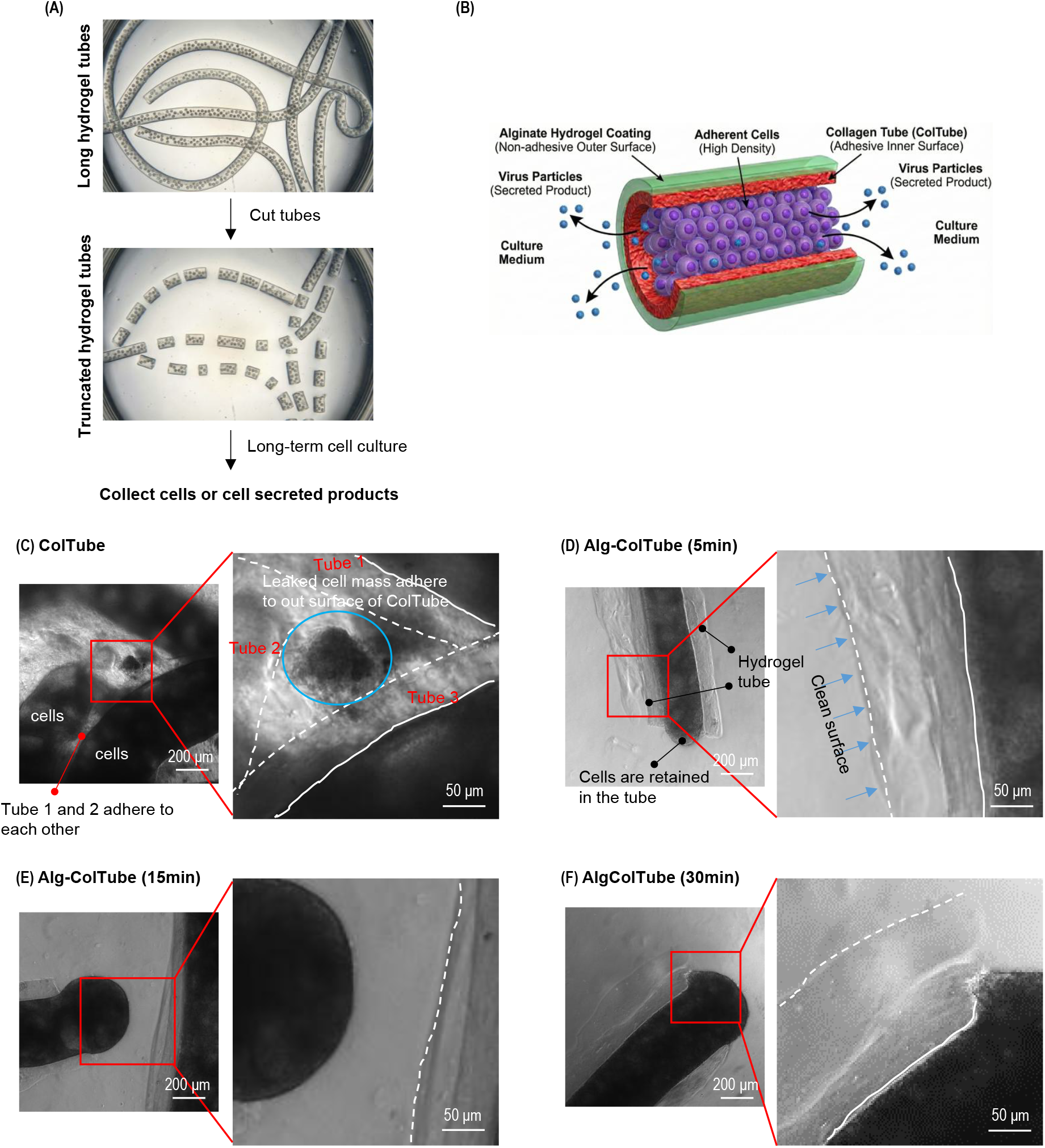
Truncated AlgColTubes for cell culture and product release. Cells are first processed into long ColTubes, which are then coated with an alginate hydrogel. These AlgColTubes support high-density cell growth. After culture, the tubes are cut into short segments, exposing both ends to the surrounding medium (A). This configuration allows large proteins and particles—such as extracellular vesicles and viruses—to be released through the open ends (B). Without alginate coating, truncated ColTubes tend to adhere to each other. Additionally, some cells leak out, attach to the outer surface, and promote tube aggregation (C). In contrast, alginate-coated tubes retain cells within the truncated segments, preventing leakage and surface adhesion (D–F). This approach enables long-term cell culture in truncated AlgColTubes while facilitating continuous collection of cell-secreted products. The technology is particularly useful for producing bioparticles such as viruses and extracellular vesicles. Solid and dashed white lines denote the inner and outer walls of Coltube or AlgColTube, respectively.

This approach exploits the spatially differentiated surface properties of AlgColTubes: cells adhere to and grow on the adhesive collagen inner surface, while the non-adhesive alginate exterior prevents tube-tube aggregation and cell growing on the outer surfaces (Fig. 8D–F). In contrast, truncated uncoated ColTubes aggregate upon cutting, with leaked cells bridging adjacent segments (Fig. 8B, C). AlgColTube segments remained dispersed and cells were retained within the lumen with no observable outer-surface fouling, enabling stable long-term culture in the truncated format.

To validate this approach for viral vector manufacturing, HEK 293SF-LVP #3E9 cells — engineered for cumate-inducible lentivirus expression — were cultured in ColTubes (Fig. 9A) or AlgColTubes (Fig. 9B) for 5 days, induced for lentivirus production, and the tubes were cut into 3-cm, 1-cm, and 1-mm segments. Cells grew to high density (~3.0×10^9^ cells/mL) with excellent viability in both formats. As expected, truncated ColTube segments adhered to each other and to the culture plate; AlgColTube segments showed no inter-tube or tube-vessel adhesion, and cells remained retained within the lumen after cutting. Lentivirus titer measurements in conditioned medium at 72 h post-induction showed that approximately 20%, 40%, and 100% of total produced virus was released for 3-cm, 1-cm, and 1-mm segments, respectively (Fig. 9C), with each shorter group significantly exceeding the next longer group (** p<0.001 vs. intact; ### p<0.001 vs. 3-cm). There were no significant differences in release percentage between ColTubes and AlgColTubes of the same segment length, indicating that segment length — not coating identity — governs release efficiency. These results establish truncated AlgColTubes as a practical platform for scalable, continuous manufacturing of biotherapeutic particles including viral vectors and extracellular vesicles.

**Fig. 9.**
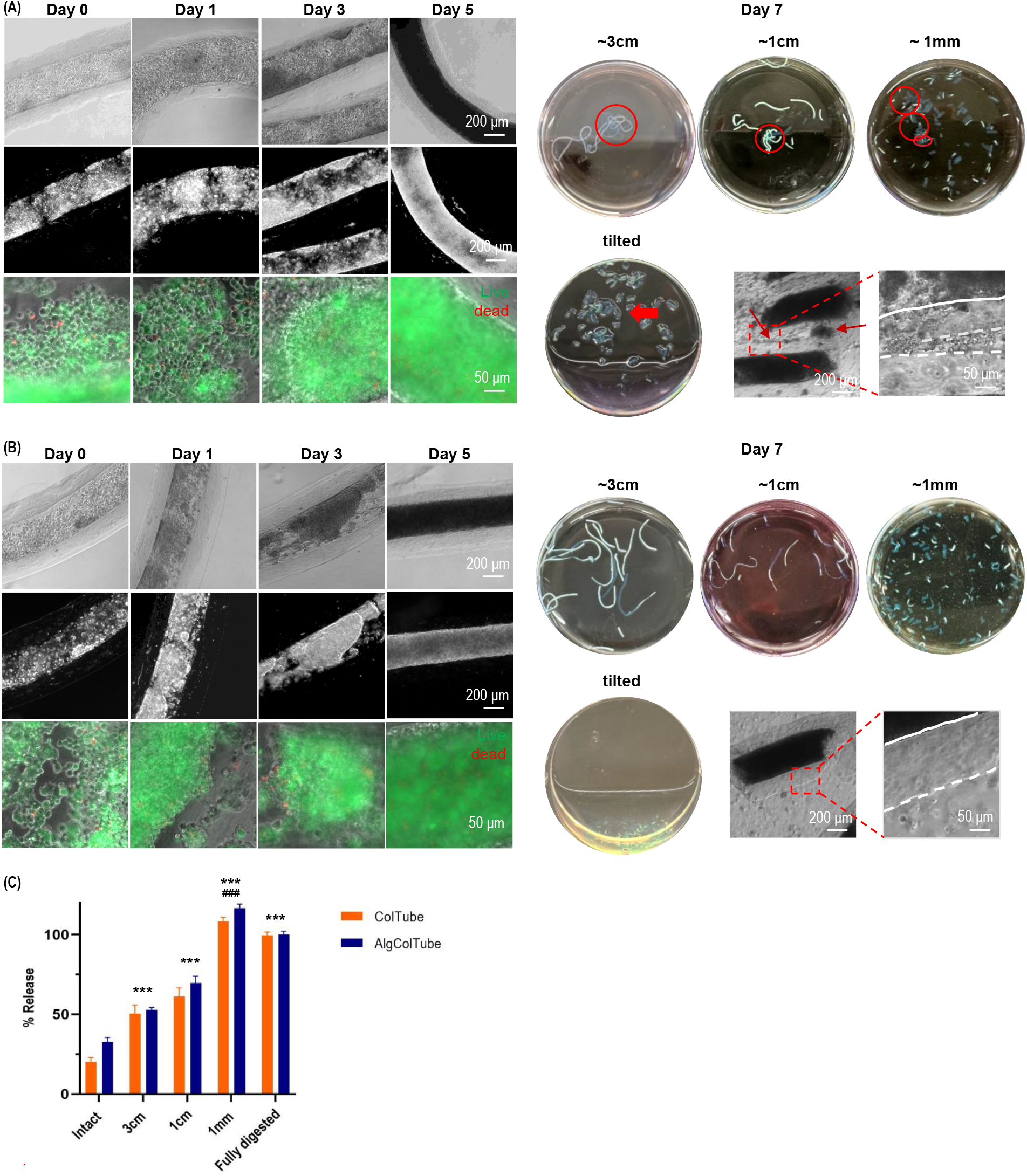
Lentivirus production using HEK 293SF-LVP #3E9 cells in truncated AlgColTubes. HEK 293SF-LVP #3E9 cells were cultured in ColTubes (A) or AlgColTubes (B). Cells grew well in both formats. After 5 days of culture, the cells were induced to produce lentivirus and the tubes were truncated into 3-cm, 1-cm, and 1-mm segments. Truncated ColTubes stuck to neighboring tubes (red circle), when AlgColTubes did not. Upon tilting, AlgColTubes settled freely, while ColTubes commonly adhered to the plate bottom (red arrow). The percentage of virus released into the medium (C) were significantly higher in truncated AlgColTubes and increased progressively as segment length decreased, indicating that open ends facilitate viral particle release into the medium. ** p < 0.001 vs. Intact group; ### p < 0.001 vs. 3cm group.

## D. Discussion

Mammalian cells underpin an expanding landscape of therapeutic and research applications that collectively demand production at extraordinary scale. Human pluripotent stem cells (hPSCs) — encompassing both human embryonic stem cells (hESCs) and induced pluripotent stem cells (iPSCs) — and their differentiated progenies are essential for treating degenerative diseases, modeling pathological states, and enabling high-throughput drug discovery16. Immune cells, including T cells and natural killer (NK) cells, are increasingly deployed in cancer immunotherapies such as CAR-T and CAR-NK cell therapies17–21. The cell quantities required for even a single patient are immense: approximately 10^9^ cardiomyocytes or β cells are needed per patient for myocardial infarction or Type 1 diabetes therapy, respectively, and roughly 10^5^ surviving dopaminergic neurons are required for Parkinson’s disease 22. When multiplied across the millions of affected individuals — over 8 million in the US living with myocardial infarction, 1–2.5 million with Type 1 diabetes, and over 1 million with Parkinson’s disease — the aggregate demand is at a scale no current manufacturing platform can sustainably meet22. The demand for scalable biomanufacturing extends equally to biotherapeutic particles — a rapidly growing class of biologics that includes viral vectors (AAV, lentivirus), virus-like particles (VLPs), and extracellular vesicles (EVs) and exosomes23–25. Scalable, consistent, and cost-effective production of all these particle types remains a critical unmet need in translational and industrial bioproduction 16,22,26.

Despite decades of bioprocess engineering development, current cell culture platforms fall substantially short of the yield, consistency, and scalability these applications require. Two-dimensional flask-based systems are intrinsically limited by their planar geometry, making them impractical for any application requiring large numbers of cells1–3,16,26. Three-dimensional suspension cultures in stirred-tank bioreactors were developed to overcome these constraints but introduce a distinct and difficult set of problems. In adherent and semi-adherent cell types including hPSCs, uncontrolled aggregation produces cell clusters that rapidly exceed the ~400 µm diffusion limit, generating hypoxic, nutrient-depleted cores that drive apoptosis, suppress proliferation, and trigger undesired differentiation 22,27–29. The agitation required to break up aggregates and improve mass transport simultaneously exposes cells to shear stress that is damaging at the intensities needed for effective mixing 22,30,31. The net result is that hPSCs in stirred-tank bioreactors typically achieve less than 10-fold expansion over four days, with volumetric yields of approximately 2×10^6^ cells/mL — utilizing less than 0.5% of available bioreactor volume32–34.

The hydrodynamic environment in stirred-tank bioreactors is also notoriously difficult to control and reproduce: shear stress profiles, chemical gradients, and aggregate size distributions all vary with bioreactor geometry, agitation speed, and medium rheology, and these parameters are scale-dependent 16,22,26,30–35. The production difficulty is illustrated in a cardiomyocyte differentiation study in ~100 mL bioreactors, where total cell yield and cardiomyocyte purity varied by more than two-fold across replicate batches using the same cell line, and changed substantially when a different hPSC line was used under identical conditions36,37. Scaling from ~100 mL to ~1000 mL would require complete re-optimization of bioreactor design, agitation rates, and differentiation protocols — a technically and economically prohibitive undertaking for most applications 36,37. The largest reported hPSC suspension cultures remain confined to tens of liters22,38, far below what therapeutic applications at population scale would require.

The hydrogel tube microbioreactor platform directly addresses these manufacturing challenges by physically decoupling mass transport efficiency from hydrodynamic shear. AlgTubes — the first generation of this platform — established that microscale hydrogel tube encapsulation enables volumetric yields of ~5×10^8^ cells/mL in hPSC cultures, more than 200-fold beyond stirred-tank systems, and supports successful culture across multiple cell types including endothelial cells, neural stem cells, vascular smooth muscle cells, and T cells ^4–13^. However, because alginate is not a native ECM component of human tissue, AlgTubes cannot fully replicate in vivo stem cell niches — a limitation relevant for ECM-sensitive cells ^14^. ColTubes addressed this gap by replacing alginate with type I collagen, providing native integrin-binding sites and fibrillar nanoarchitecture ^14^. Yet collagen’s biological adhesivity — while productive at the inner surface — created an outer-surface liability that would preclude scale-up. AlgColTubes resolve this fundamental tension by engineering spatially differentiated surface properties: a cell-instructive collagen interior and a non-adhesive alginate exterior.

The durability of the alginate coating under both static and dynamic conditions is mechanistically grounded in IPN formation rather than simple surface adsorption. Alginate penetrates the hydrated collagen matrix during the dipping step and becomes physically entangled with collagen nanofibers before Ca^2+^-mediated ionic crosslinking locks the network in place. The resulting IPN resists delamination that would be expected for a purely surface-adsorbed layer. SEM data show that penetration depth scales with alginate concentration and dipping time (Figs. 2, 3). Importantly, because the IPN is heterogeneous — denser at the outer surface, transitioning to native collagen nanofibers at the inner surface — it preserves the cell-adhesive character of the collagen interior while presenting an anti-fouling alginate face externally. The concentration and time of coating can thus serve as independent design parameters to balance anti-adhesion performance, wall diffusivity, and ECM presentation for specific cell types or applications.

Alginate’s anti-adhesive surface properties arise from its hydrophilic, negatively charged character: the dense hydration shell sterically excludes approaching proteins and cells, while electrostatic repulsion further disfavors adhesion of cells carrying net-negative surface charges. At coating times of ≥15 min, even a thin but continuous alginate surface layer is sufficient to completely suppress all measured adhesion — tube-tube, tube-vessel, and outer-surface cell fouling — under both static and 60-rpm dynamic conditions. This has direct and important implications for bioreactor scale-up, where hundreds of milliliters to liters of tube suspension must be maintained as a free-flowing, non-aggregating dispersion. Adhesion events in large-scale culture would cause irreversible clumping, create diffusion-limited micro-zones, and complicate suspension handling during medium exchange and harvest. AlgColTubes eliminate all of these risks at a coating time that is rapid, reagent-inexpensive, and readily integrated into standard ColTube fabrication workflows.

The ability to fabricate stable, non-aggregating truncated tube segments — retaining cells internally while releasing particles through open ends — is a functionally novel capability enabled specifically by the anti-adhesive outer surface of AlgColTubes. The near-complete (~100%) lentivirus release from 1-mm AlgColTube segments (Fig. 9C) indicates that the tube wall is the primary barrier to particle egress in intact tubes, and that reducing segment length below a critical threshold effectively eliminates this barrier. Future studies should characterize release kinetics for other particle types — including AAV, EVs, and exosomes — across a broader range of particle sizes and segment lengths, and should explore whether perfusion-based delivery of fresh medium through truncated AlgColTube arrays could enable true continuous manufacturing formats relevant to clinical gene therapy vector production, where batch yield variability remains a significant challenge23,25.

## Author Contributions

Y.L. conceived the idea and directed the work. Y.L., X.W., and Y.Y. designed the experiments. Y.Y., X.W., Y.P., and L.H. performed experiments. A.M. provided cells. Y.L., X.W., Y.Y., A.M., Y.W., and L.H. analyzed the data and wrote the manuscript.

## Data Availability

The authors confirm that the data supporting the findings of this study are available within the article.

## Ethical Statement

All experiments involving human embryonic stem cells (hESCs) were performed in strict accordance with institutional guidelines and approved by the appropriate ethics committee at Pennsylvania State University. The hESC line used in this study was H9, obtained from the WiCell Research Institute and authorized for research use. No new hESC derivation was conducted. All procedures complied with applicable national regulations and policies governing the ethical use of human stem cells in research.

