## Supplemental figure 1 for "Alginate-Coated Collagen Hydrogel Tubes for Scalable Cell and Biotherapeutic Particle Manufacturing"

### Slide 1
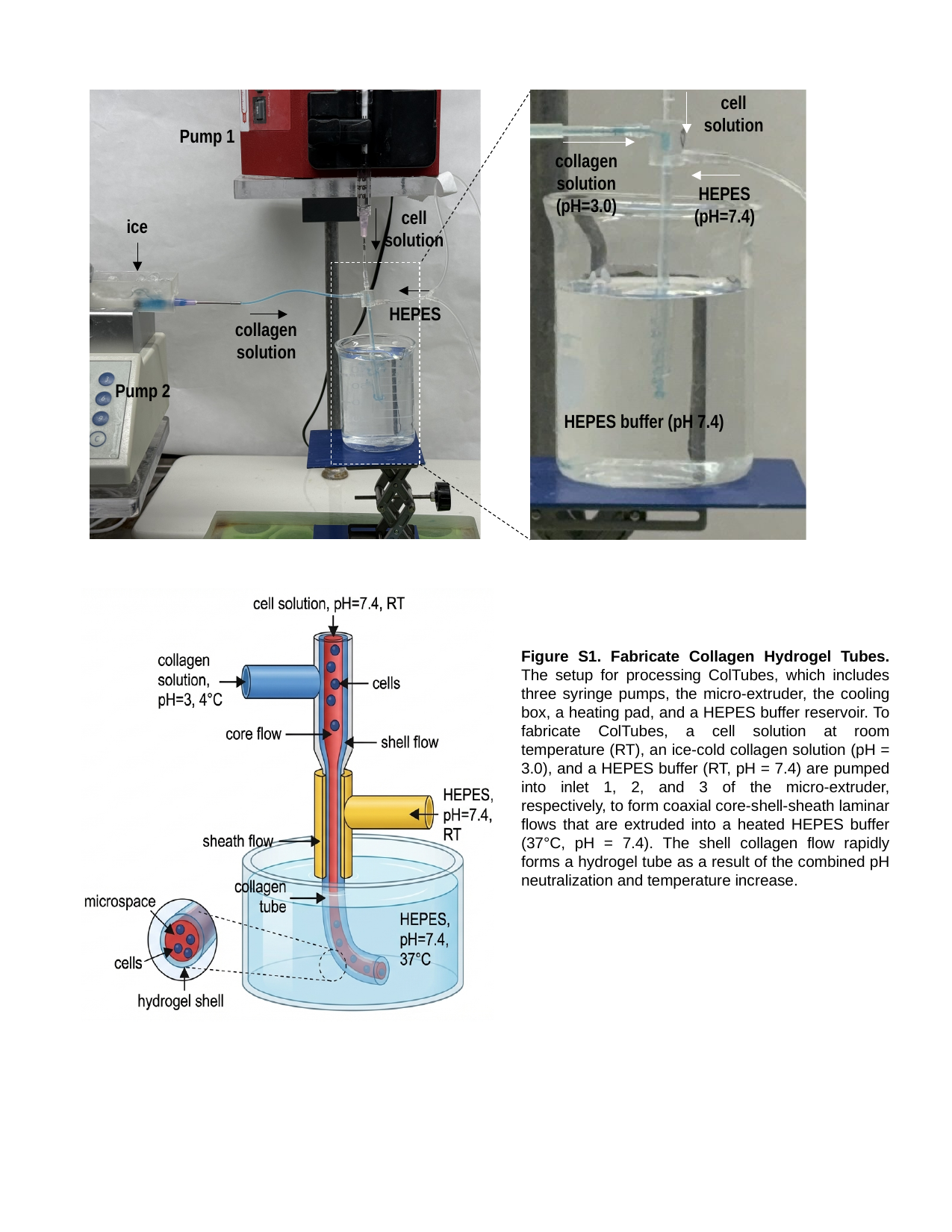

cell solution
Pump 1
collagen solution (pH=3.0)
HEPES (pH=7.4)
cell solution
ice
HEPES
collagen solution
Pump 2
HEPES buffer (pH 7.4)
Figure S1. Fabricate Collagen Hydrogel Tubes. The setup for processing ColTubes, which includes three syringe pumps, the micro-extruder, the cooling box, a heating pad, and a HEPES buffer reservoir. To fabricate ColTubes, a cell solution at room temperature (RT), an ice-cold collagen solution (pH = 3.0), and a HEPES buffer (RT, pH = 7.4) are pumped into inlet 1, 2, and 3 of the micro-extruder, respectively, to form coaxial core-shell-sheath laminar flows that are extruded into a heated HEPES buffer (37°C, pH = 7.4). The shell collagen flow rapidly forms a hydrogel tube as a result of the combined pH neutralization and temperature increase.
